# Sentinel versus passive surveillance for measuring changes in dengue incidence: Evidence from three concurrent surveillance systems in Iquitos, Peru

**DOI:** 10.1101/040220

**Authors:** Sandra Olkowski, Steven T. Stoddard, Eric S. Halsey, Amy C. Morrison, Christopher M. Barker, Thomas W. Scott

**Affiliations:** Department of Entomology and Nematology, University of California, Davis, CA, USA; U.S. Naval Medical Research Unit No. 6, Lima, Peru; Department of Pathology, Microbiology and Immunology, University of California, Davis, CA, USA; Fogarty International Center, National Institutes of Health, Bethesda, MD, USA

**Author notes:** Corresponding author: Sandra Olkowski.

**Keywords:** dengue, epidemiology, surveillance, cohort, sentinel surveillance

## Abstract

Monitoring changes in infectious disease incidence is fundamental to outbreak detection and response, intervention outcome monitoring, and identifying environmental correlates of transmission. In the case of dengue, little is known about how consistently surveillance data track disease burden in a population over time. Here we use four years of monthly dengue incidence data from three sources – population-based (‘passive’) surveillance including suspected cases, ‘sentinel’ surveillance with 100% laboratory confirmation and complete reporting, and door-to-door (‘cohort’) surveillance conducted three times per week - in Iquitos, Peru, to quantify their relative consistency and timeliness. Data consistency was evaluated using annual and monthly expansion factors (EFs) as cohort incidence divided by incidence in each surveillance system, to assess their reliability for estimating disease burden (annual) and monitoring disease trends (monthly). Annually, passive surveillance data more closely estimated cohort incidence (average annual EF=5) than did data from sentinel surveillance (average annual EF=19). Monthly passive surveillance data generally were more consistent (ratio of sentinel/passive EF standard deviations=2.2) but overestimated incidence in 26% (11/43) of months, most often during the second half of the annual high season as dengue incidence typically wanes from its annual peak. Increases in sentinel surveillance incidence were correlated temporally (correlation coefficient = 0.86) with increases in the cohort, while passive surveillance data were significantly correlated at both zero-lag and a one-month lag (0.63 and 0.44, respectively). Together these results suggest that, rather than relying on a single data stream, a clearer picture of changes in infectious disease incidence might be achieved by combining the timeliness of sentinel surveillance with the representativeness of passive surveillance.

## Introduction

Infectious disease surveillance in developing countries is often challenged by limited public health resources, insufficient laboratory capacity, and incomplete reporting [1]. In order to obtain high-quality data in the face of these and other challenges, the World Health Organization (WHO) has recommended sentinel surveillance for many infectious diseases [2,3]. In sentinel surveillance systems, resources are focused on collecting complete, timely data from a subset of healthcare facilities or laboratories [4], thus requiring fewer resources than would be needed to actively collect the same quality of data from all facilities (population-based active surveillance). Passive surveillance systems, in which data collection is dependent on reporting by healthcare facilities, are representative by virtue of being population-based, but are also subject to under-detection and underreporting [5]. The goal of this study is to evaluate the public health utility of sentinel surveillance compared to passive surveillance for measuring changes in an endemic infectious disease, using dengue as a case study.

While it is well-established that data from passive surveillance underestimate the incidence (in this study: the number of new symptomatic cases per 100,000 persons per month) of infectious diseases [5] including dengue [6], the nature of temporal variation in underdetection is less clear. Such variation could have significant public health implications if the meaning of surveillance-based incidence changes over time. Here we express variation using a monthly expansion factor (EF) (i.e., the ratio of incidence in cohort surveillance to incidence in sentinel or passive surveillance). This method has been used in previous dengue studies [7,8] to estimate annual disease burden, but there is no published evidence regarding the consistency of inter-annual EFs are and thus how effectively they might be applied to estimating finer-scale changes in dengue incidence.

Although laboratory-based sentinel surveillance has been recommended for dengue [9], the WHO urges caution in over-interpreting these data [10] because sentinel surveillance may not adequately represent broader population trends in incidence. Assessing the added value of sentinel surveillance over passive surveillance for capturing a consistent proportion of cases and detecting seasonal increases in incidence would require that both be compared against data from a third ‘gold-standard’ system that provides an objective baseline measure of incidence [11]. Thus we compared data from both systems to reference data gathered from community-based surveillance in a longitudinal cohort.

Dengue is an acute febrile illness (AFI) caused by any one of four serotypes of the dengue virus (DENV). It is the most prevalent mosquito-borne virus globally and is a growing health concern, with an estimated incidence of 96 million symptomatic infections per year [12]. Here, dengue incidence measurement is considered in the context of Iquitos, Peru, where the disease most commonly presents as an undifferentiated, self-limiting AFI with an annual high transmission season. This scenario highlights two challenges that could potentially be addressed by sentinel surveillance data. First, because it is often clinically undifferentiated, surveillance data that include suspected cases will be influenced by physicians’ assumptions about transmission. These could potentially be improved with laboratory-confirmed sentinel surveillance, which would provide a more objective measure of the proportion of DENV cases among febrile individuals who seek treatment. Second, timely indication of increased incidence is of special concern in dengue because reactive mosquito-control activities are a common public health intervention. Rapid identification of changes in incidence based on sentinel surveillance data may result in a more effective response.

In Iquitos, there are three concurrent dengue surveillance systems: passive, sentinel, and door-to-door febrile surveillance in a longitudinal research cohort. Here we use cohort data as a reference to test the hypothesis that sentinel surveillance data are a more consistent – thus more ‘accurate’ – measure of incidence by month and also a more timely indicator of seasonal increases in incidence than passive surveillance data.

## Methods

### Study area and study designs

Iquitos is a geographically isolated city of ~440,000 inhabitants, located in the Amazon basin of northeastern Peru. DENV was re-introduced into the city in 1990 and was a reportable disease during our study. Beginning in 1990, each DENV serotype has been introduced and subsequently dominated transmission for multiple years before being replaced [13]. The current study includes data from three sources, detailed below: (1) ‘passive surveillance’ (confirmed and suspected cases reported by all healthcare facilities to the Directión Regional de Salud Loreto (DRSL)), (2) sentinel surveillance (laboratory-confirmed cases from a city-wide AFI research network), and (3) door-to-door febrile surveillance in longitudinal cohorts (‘cohort surveillance’).

### Ethics statement

The de-identified data used in this study were collected under four protocols (NMRCD2000.0006, NMRCD2010.2010, NMRCD2005.009, NMRCD2007.0007), all approved by the Institutional Review Boards (IRBs) of the Naval Medical Research Center and Naval Medical Research Unit No. 6 (NAMRU-6, formerly Naval Medical Research Center Detachment). The sentinel surveillance protocol (NMRCD 2010.2010) was also approved by the Ethics Committee for the Peruvian National Institute of Health (INS, acronym in Spanish). One cohort protocol (NMRCD2005.0009) was also approved by IRBs at the University of California, Davis (UC Davis), Universidad Peruana Cayetano Heredia University in Lima, Peru, and the second (NMRCD2007.0007) received local approval from NAMRU-6 also registered as a Peruvian IRB, as well as the UC Davis IRB. All protocols were in compliance with regulations in the United States and Peru governing the protection of human subjects.

### Cohort surveillance

Cohort surveillance data are from two spatially and temporally overlapping longitudinal cohorts. One cohort was restricted to two neighborhoods and the other cohort was distributed across the northern portion of the city. There were an average of 4,700 people under febrile surveillance during the study period. In both cohorts, phlebotomists visited participating houses three times per week to monitor all individuals ≥ 5 years of age for dengue-like illness. Inclusion criteria were occurrence of fever (≥ 38°C) for ≤ five days, either by axial measurement or subject-reported in combination with the use of anti-pyretics, plus at least one other symptom consistent with DENV infection, including headache, rash, or retro-orbital pain. Positive cases were defined by DENV RNA detection by reverse transcription polymerase chain reaction (RT‐PCR) or a ≥ four-fold increase in DENV antibodies between acute and convalescent samples, as measured by IgM capture enzyme-linked immunosorbent assay (ELISA). RT-PCR and ELISA protocols were as previously described [14].

### Sentinel surveillance

Sentinel febrile surveillance was carried out by NAMRU-6. Here we include dengue case counts from two public hospitals and eight public outpatient clinics located throughout Iquitos, together serving ~208,000 residents (~47% of the population during the study period) (DRSL, unpub. data). A febrile case was defined as an individual ≥ 5 years old experiencing a fever of ≥ 38°C for a maximum of five days. These individuals were invited to participate, regardless of clinical diagnosis. An acute blood sample was collected at the time of enrollment and a convalescent sample was collected two to four weeks later, when possible. Criteria for positive cases were the same as for the cohorts, as described above, and infection detection protocols were as previously described [15].

### Passive surveillance

Passive surveillance data are based on case counts reported to the DRSL. These include suspected and laboratory-confirmed dengue cases from the four districts that comprise the city of Iquitos, located in the Department of Loreto. Iquitos is estimated to account for ~64% of all dengue cases in the region [16]. The total number of reported cases was scaled accordingly. Case data were restricted to individuals ≥ 5 years of age, to correspond with the cohort and sentinel surveillance study protocols.

### Study period characteristics

Two successive DENV introductions occurred during the study period of 1 July 2008 to 30 June 2012. A virgin-soil invasion of dengue virus 4 (DENV-4) occurred in October 2008 [17] and a novel genotype of DENV-2 American/Asian lineage II (DENV-2) was introduced in November 2010 [18]. Both introductions resulted in replacement events, so that by the second year of circulation (‘interepidemic’ years), the introduced virus accounted for ≥ 90% symptomatic DENV infections identified in both cohort and health-center based surveillance.

Based on ten years of data, high dengue incidence in Iquitos was observed between September and April; peak incidence occurred at various points in that interval. We used trimester transmission season designations: (1) low, May to August, (2) early high, September to December, and (3) late high, January to April [12]. While these designations do not reflect observed incidence pattern for every year, they delineate periods of transmission as perceived by patients, physicians, and public health officials.

### Analyses

All analyses were conducted by month (July to June, annually) to minimize the number of time periods with zero in the denominator of the incidence rate ratio (i.e., sentinel surveillance/cohort surveillance). Out of a possible 48 months, 43 were included in the analysis of passive surveillance data. Five months were excluded because no dengue cases were identified in the cohorts. Analysis using sentinel surveillance excluded two additional months: one because no cases were identified by sentinel surveillance and another when the participants were not enrolled for most of the month. Null months were distributed across transmission season categories, with three in the low season and two in each of the other seasons.

Cohort incidence was defined as dengue cases per 100,000 persons per month. The surveillance population included all persons residing in houses monitored three times per week by door-to-door febrile surveillance in the longitudinal cohorts. Sentinel surveillance incidence was defined as dengue cases per 100,000 persons per month in the combined catchment populations of participating hospitals and clinics. Catchment areas were estimated in 2008 by the DRSL (unpub. data). Passive surveillance incidence included both suspected and confirmed dengue cases per 100,000 persons per month in the total estimated 2008 population of Iquitos [19]. To match the restrictions in cohort case data, population estimates used to calculate incidence for sentinel and passive surveillance were restricted to persons ≥5 years of age. All individuals under surveillance by any of the three methods were assumed to contribute person-time to the incidence estimate.

We calculated annual and monthly EFs for passive and sentinel surveillance to describe the range of healthcare based case detection, relative to cohort surveillance, within each year and between years. EFs were calculated as cohort incidence divided by either passive or sentinel incidence. Ratio of standard deviations was used to compare variation in data by year and season.

To compare the relative timing of passive versus sentinel surveillance systems for early identification of seasonal increases in dengue incidence, we performed a cross-correlation analysis of both systems with cohort surveillance for the full time series and by transmission season. This method quantifies the strength of the temporal association between the cohort surveillance incidence rate at month *t* and the passive or sentinel surveillance incidence rate at month *t+h* (where the sign of *h* indicates a temporal lag or lead).

Statistical analysis was performed using R version 3.1.1 [20]. Statistical significance was assessed at α = 0.05.

## Results

### Incidence

In each of the four transmission years (July to June) studied, the highest annual and seasonal incidence occurred in cohort surveillance, followed by passive surveillance, and the lowest incidence occurred in sentinel surveillance (Figure 1, Table 1). An exception was the DENV-2 introduction (2010-11), when passive surveillance case numbers surpassed cohort surveillance in the late high season (January-April).

**Figure 1:**
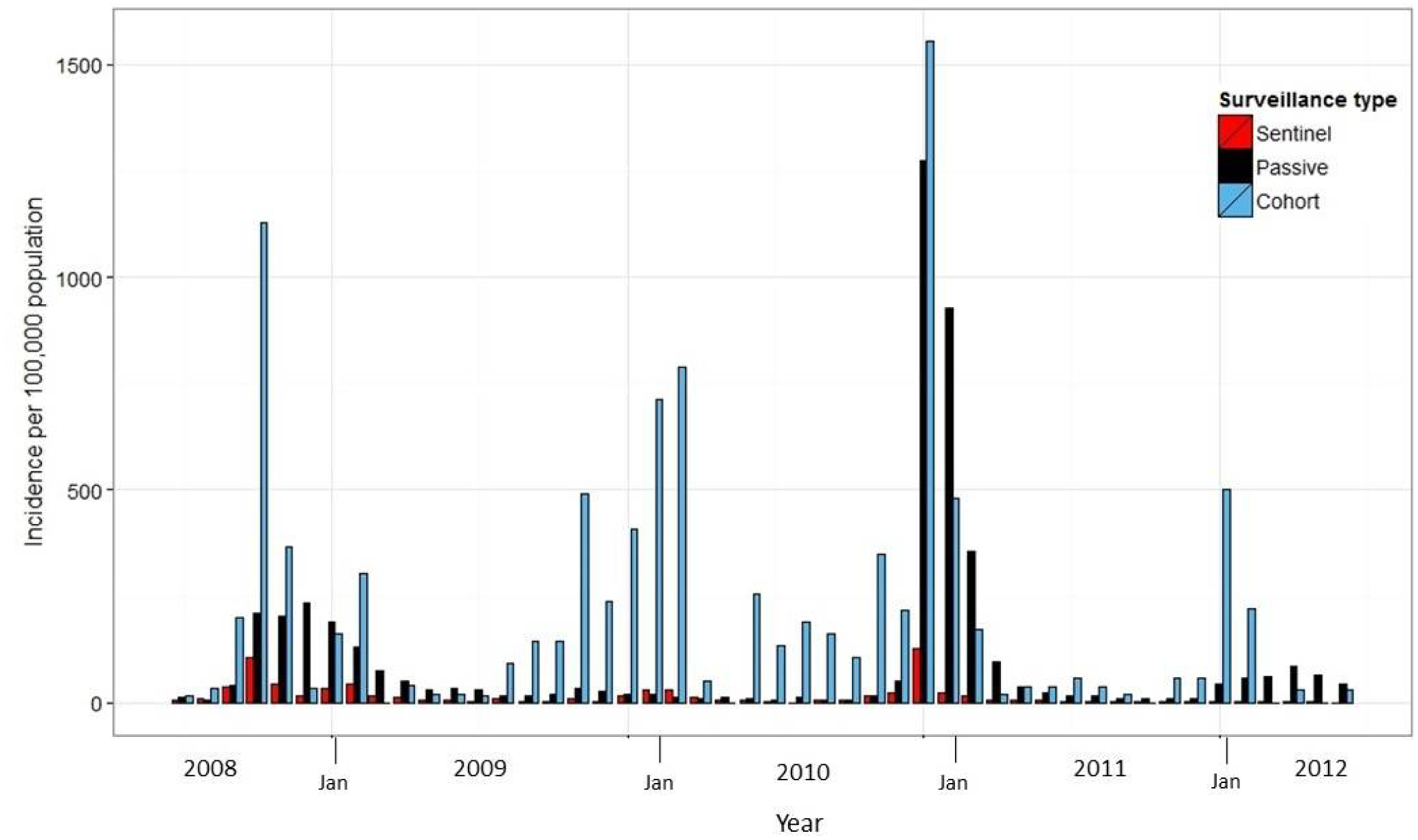
Incidence measured as per 100,000 population per month by sentinel, passive, and cohort surveillance. Bars are grouped by month to represent incidence as measured by each of the three surveillance systems.

**Table 1:** Incidence per 100,000 persons per time period, and associated expansion factors (EFs).

|  | Cohort surveillance | Sentinel surveillance | Passive surveillance |
| --- | --- | --- | --- |
| **2008-09** |  |  |  |
| Annual total | 2324.6 | 339.9 | 1220.4 |
| Low season total | 90.5 | 27.6 | 82.1 |
| Early high season total | 1725.3 | 204.6 | 691.3 |
| Late high season total | 508.8 | 107.7 | 447.1 |
| Monthly range<sup>b</sup> | 16.6 – 1128.1 | 5.3 – 106.9 | 4.6 – 233.3 |
| Peak incidence month | October | October | December |
| Annual EF | NA | 6.8 | 1.9 |
| Monthly EF range | NA | 2.1 – 10.6 | 0.14 – 7.2 |
| **2009-10** |  |  |  |
| Annual total | 3341.4 | 133.5 | 236.08 |
| Low season total | 363.5 | 25.0 | 72.7 |
| Early high season total | 1020.5 | 19.2 | 97.5 |
| Late high season total | 1957.4 | 89.3 | 65.9 |
| Monthly range<sup>a</sup> | 18.2 – 788.0 | 1.4 – 29.8 | 11.2 – 33.2 |
| Peak incidence month | March | February | November |
| Annual EF | NA | 25.0 | 14.2 |
| Monthly EF range | NA | 4.2 – 75.9 | 0.60 – 55.2 |
| **2010-11** |  |  |  |
| Annual total | 3461.8 | 244.0 | 2810.0 |
| Low season total | 400.4 | 13.9 | 81.5 |
| Early high season total | 836.0 | 51.9 | 76.5 |
| Late high season total | 2225.4 | 177.7 | 2651.9 |
| Monthly range<sup>b</sup> | 19.2 – 1554.0 | 1.4 – 128.7 | 4.6 – 1272.8 |
| Peak incidence month | January | January | January |
| Annual EF | NA | 14.2 | 1.2 |
| Monthly EF range | NA | 2.5 – 93.6 | 0.20 – 31.4 |
| **2011-12** |  |  |  |
| Annual total | 1015.8 | 33.1 | 427.3 |
| Low season total | 128.0 | 6.2 | 141.0 |
| Early high season total | 135.3 | 12.0 | 38.6 |
| Late high season total | 752.5 | 14.9 | 247.7 |
| Monthly range<sup>b</sup> | 19.3 – 501.7 | 1.4 – 4.3 | 8.8 – 120.7 |
| Peak incidence month | January | December-March<sup>b</sup> | April |
| Annual EF | **NA** | 30.7 | 2.4 |
| Monthly EF range | **NA** | 13.4 – 116.1 | 0.4 – 11.3 |

<sup>a</sup> Does not include months in which no cases were detected
<sup>b</sup> Equal incidence across four months

At least 80% of cases were identified between September and April in each surveillance system. Peak incidence generally occurred in the early high season (September-December) or early in the late high season (January-April) (Figure 1, Table 1).

### Time-varying incidence

Annual EFs from 2008-2012 based on passive surveillance relative to cohort surveillance were consistently lower than those based on sentinel surveillance data: 1.9 vs. 6.8, 14.2 vs. 25.0, 1.2 vs 14.2, 2.4 vs. 30.7 (Table 1). Both data series were highly variable by month, compared to annual figures (Figure 2, Table 1). The overall relationship of sentinel surveillance data to cohort data was more variable than that of passive surveillance data to cohort data, as measured by the relative standard deviation of monthly EFs (ratio of sentinel/passive EF standard deviations = 2.2). This pattern was also observed by year (ratio of standard deviations = 1.1, 1.4, 2.2, 10.8) and season (ratio of standard deviations = 13.2, 2.6, 1.8 for low, early high, and late high seasons, respectively).

**Figure 2:**
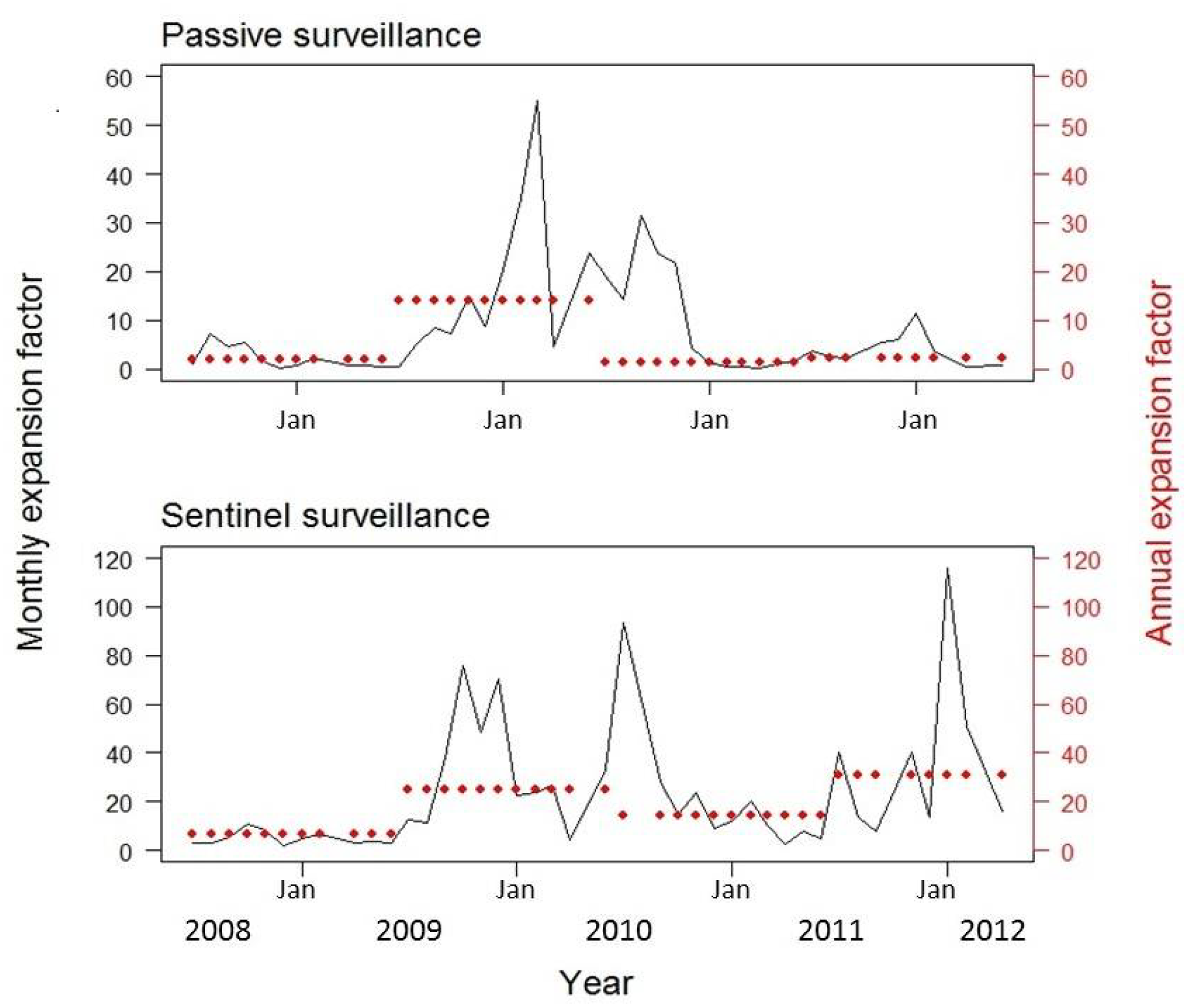
Monthly and annual expansion factors calculated as the ratio of cohort to passive (top panel) and sentinel (bottom panel) incidence (monthly or annual cases per 100,000 population).

Passive surveillance overestimated dengue incidence in 26% (11/43) of months (indicated by EF values <1). Low (May-August), early high (September-December), and late high (January-April) seasons contained four, one and six of these overestimated months, respectively. Sentinel surveillance always underestimated incidence (i.e., EF >1).

Monthly increases in sentinel surveillance incidence were correlated with increases in cohort surveillance cases during the same month, across the time series (correlation coefficient = 0.86), as well as early (0.87) and late (0.84) high transmission seasons (Figure 3). Increases in passive surveillance incidence were also associated with increases in cohort incidence overall and by transmission season. The full time series showed statistically significant associations both in the same month and with a one-month lag (0.63 and 0.44, respectively). The strongest association in the early high season was at a one-month lag (correlation=0.57). In the late high season, incidence was highly correlated without any lag (correlation=0.65). Both systems showed mixed positive and negative correlations in the low season, none of which were significant.

**Figure 3:**
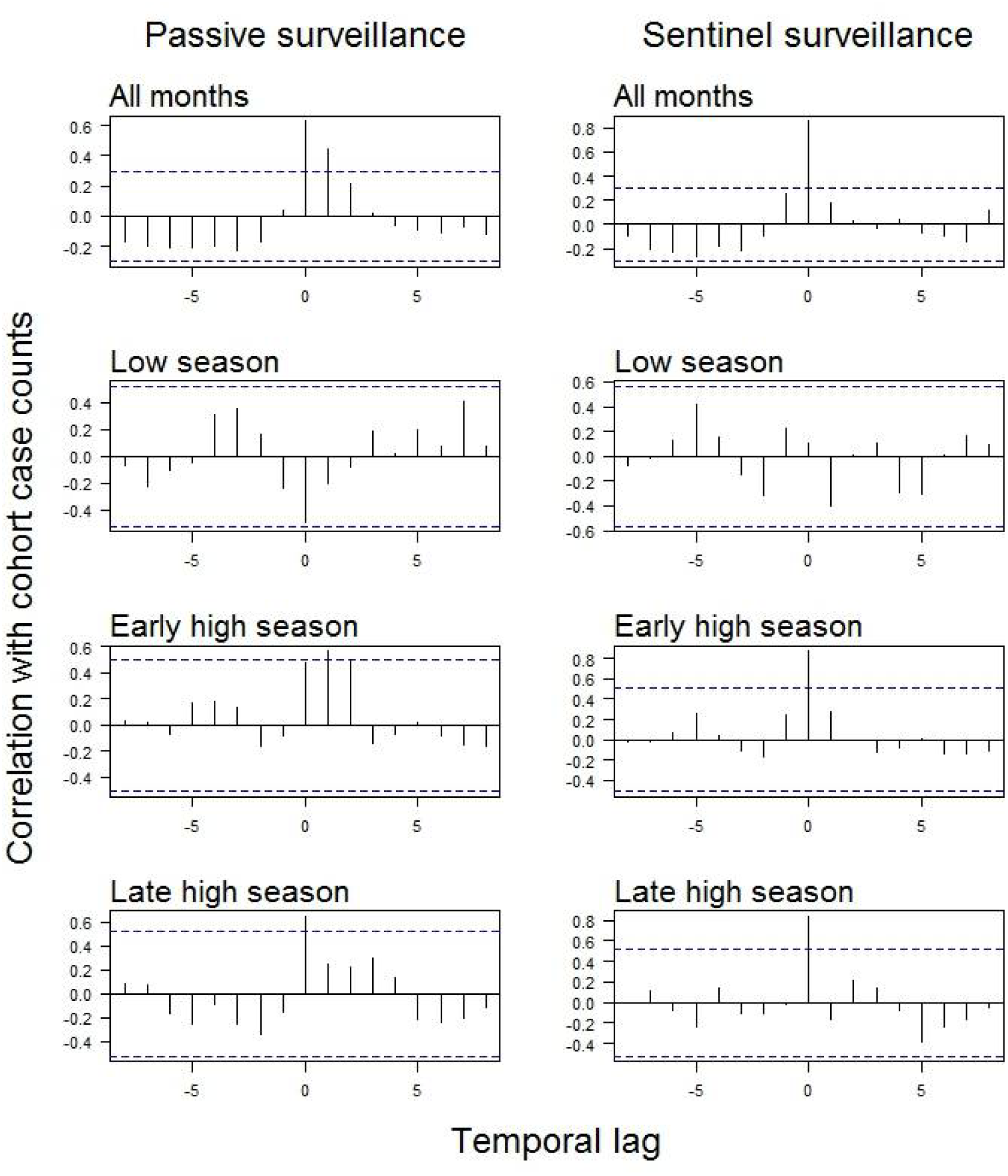
Temporal correlation of case detection for passive surveillance and sentinel surveillance relative to cohort surveillance. Positive numbers on the x-axis indicate a delayed increase in cases. Positive numbers on the y-axis indicate a positive relationship (i.e., sentinel or passive surveillance case numbers increase as active surveillance case numbers increase). Horizontal dotted lines represent statistical significance at α = 0.05

## Discussion

Here we use four years of data from three concurrent dengue surveillance systems in Iquitos, Peru, to assess the relative performance of monthly data from sentinel surveillance and passive surveillance, based on the criteria of consistency and timeliness in relation to a referent cohort incidence time series. Sentinel surveillance data generally reflected seasonal increases in dengue incidence earlier than in passive surveillance – in the same month as the cohort, as opposed to a lag of up to two months – but were not a reliable indicator of the magnitude of the increase. Data from passive surveillance, on the other hand, were generally more consistent, in that they had a lower range of EF values, but overestimated incidence in 26% (11/43) of months, most often during the second half of the annual high transmission season.

An annual EF, calculated by comparing cohort incidence to population-based surveillance data, can be used to estimate total disease burden and set public health priorities. Here, we found that passive surveillance data were a closer estimate of annual disease burden as measured using cohort data. In three of the four years, passive surveillance data included approximately half of all cases expected based on the cohort incidence, resulting in an EF of ~2. During 2009-2010, the EF rose to ~1. This change may have been driven by a lower sense of reporting urgency because DENV-4 had circulated in the previous year and was perceived to cause only mild illness. We extended the analysis to consider monthly EFs, with the goal of understanding how consistently sentinel surveillance data captures finer-scale trends when the objective is to monitor intervention outcomes or understand temporal changes in transmission patterns. Compared to passive surveillance, monthly data from the sentinel study had an overall wider range of EF values (ratio of sentinel/passive standard deviations = 2.2).

The temporal relationship between increased case counts in surveillance data and what actually happens in the population is of primary interest when the surveillance objective is to detect and rapidly respond to potential outbreaks. For infectious diseases with a seasonal pattern, such as influenza and dengue, the period of greatest interest is early in the annual high season. We found that sentinel surveillance data show a strong positive correlation to cohort data (i.e., both have rising case counts) within the same month during the trimester when increased incidence might first be expected. Conversely, the strongest correlation for passive surveillance was observed at a one-month lag, although there were similar correlations at zero and two months. Our finding that > 50% (6/11) of the months in which incidence was overestimated by passive surveillance data occurred later in the ‘expected’ high season suggests that this temporal variation in identifying increased incidence may be due to seasonal differences in case reporting and/or physicians’ index of suspicion, so that detection rates are a product of both expectation of dengue and actual incidence.

The advantage of early indication of potential outbreaks provided by sentinel surveillance data is tempered by their inconsistent proportional relationship to incidence in the community. In the early high season, the monthly EF for sentinel surveillance data varied much more than for passive surveillance data (ratio of sentinel/passive standard deviations = 2.6). Identifying increased incidence, when that might typically be expected, is necessary but likely not sufficient for triggering a costly vector control response in most dengue-endemic areas, depending on public health resources a public health department for responding to what may be a false alarm.

Observational study data only approximate ‘true’ surveillance or population incidence data. While this analysis is the first comparison of sentinel surveillance data with cohort data for dengue, the sentinel surveillance data considered here are based on hospital and clinic patients agreeing to participate in the study, rather than the total number of those seeking medical treatment. However, this sampling effect may be offset by complete data reporting in the context of a scientific study. These sentinel data might also introduce an age-related bias by consistently capturing a lower proportion of pediatric patients, based on observations by our study phlebotomists that children are generally less willing than adults to consent to a venous blood draw (unpub. data).

Another limitation is that cohort surveillance may not be measuring the same population as was sampled by the other two surveillance systems. Passive and sentinel surveillance data in Iquitos are drawn from healthcare facilities throughout the city, whereas one of the cohorts was distributed across the northern portion of the city and the other focused in two neighborhoods. However, strong temporal correlation between increased incidence in the cohort and in sentinel surveillance suggests that, at least early in the high transmission season, these data are not being significantly biased by the spatial heterogeneity of DENV transmission that has been observed in Iquitos [21].

Passive surveillance data are generally a more consistent measure of dengue incidence in the community, compared to sentinel surveillance, while sentinel surveillance data provide a more timely indicator of potential seasonal outbreaks, in the context of Iquitos. In other places where data from both types of surveillance systems are available, these findings can guide decisions about which data to use for specific public health objectives. For example, sentinel surveillance data may be useful in the context of an early outbreak warning system, but do not appear to be reliable for monitoring long-term pathogen transmission trends, due to their overall inconsistent relationship with population-based incidence. On the other hand, passive surveillance data more accurately measure overall incidence trends, yet often overestimate monthly incidence late in the transmission season when they include suspected cases, suggesting that physician suspicion and reporting are driving surveillance measures. An integrated surveillance system could trigger low-resource public health actions based on sentinel surveillance, such as alerting all healthcare facilities specifically to encourage timely reporting and increase healthcare awareness. This could reduce delays in passive surveillance data, while retaining their population-level representativeness. To improve the understanding, and thus the application, of surveillance data there is a need for objective measures of how human behavioral factors such as physician suspicion, case reporting, and treatment seeking influence measurements of incidence. Results of this study highlight the need to explicitly consider the implications of inconsistent detection when using infectious disease incidence data for outbreak detection and trend monitoring.

## Acknowledgements

The authors would like to thank all the study participants in Iquitos for being gracious partners in public health research. An equally invaluable contribution was made by the knowledgeable and dedicated staff who collect data in homes, clinics, and hospitals throughout Iquitos. Finally, thank you to Alex Perkins and Robert Reiner for their insightful feedback at every stage of this project.

## Funding statement

National Institutes of Health (R01 AI069341 and P01 AI098670), Armed Forces Health Surveillance Center’s Global Emerging Infections Systems Research Program, Military Infectious Disease Research Program, Research and Policy for Infectious Disease Dynamics (RAPIDD) program of the Science and Technology Directory, Department of Homeland Security, U.S. National Aeronautics and Space Administration (#NNX15AF36G), and Fogarty International Center, National Institutes of Health.

## Disclaimers

The views expressed in this article are those of the authors and do not necessarily reflect the official policy or position of the Department of the Navy, the Department of Defense, or the US government.

Author Eric S. Halsey is a military service member. This work was prepared as part of his official duties. Title 17 U.S.C. § 105 provides that ‘Copyright protection under this title is not available for any work of the United States Government’. Title 17 U.S.C. § 101 defines a U.S. Government work as a work prepared by a military service members or employees of the U.S. Government as part of those person’s official duties.

